# Gamete simulation improves polygenic transmission disequilibrium analysis

**DOI:** 10.1101/2020.10.26.355602

**Authors:** Jiawen Chen, Jing You, Zijie Zhao, Zheng Ni, Kunling Huang, Yuchang Wu, Jason M. Fletcher, Qiongshi Lu

## Abstract

Polygenic risk scores (PRS) derived from summary statistics of genome-wide association studies (GWAS) have enjoyed great popularity in human genetics research. Applied to population cohorts, PRS can effectively stratify individuals by risk group and has promising applications in early diagnosis and clinical intervention. However, our understanding of within-family polygenic risk is incomplete, in part because the small samples per family significantly limits power. Here, to address this challenge, we introduce ORIGAMI, a computational framework that uses parental genotype data to simulate offspring genomes. ORIGAMI uses state-of-the-art genetic maps to simulate realistic recombination events on phased parental genomes and allows quantifying the prospective PRS variability within each family. We quantify and showcase the substantially reduced yet highly heterogeneous PRS variation within families for numerous complex traits. Further, we incorporate within-family PRS variability to improve polygenic transmission disequilibrium test (pTDT). Through simulations, we demonstrate that modeling within-family risk substantially improves the statistical power of pTDT. Applied to 7,805 trios of autism spectrum disorder (ASD) probands and healthy parents, we successfully replicated previously reported over-transmission of ASD, educational attainment, and schizophrenia risk, and identified multiple novel traits with significant transmission disequilibrium. These results provided novel etiologic insights into the shared genetic basis of various complex traits and ASD.

## Introduction

Polygenic risk scores (PRSs) derived from summary statistics of genome-wide association studies (GWASs) have enjoyed great popularity in human genetics research. Applied to a variety of complex traits and diseases, PRS can effectively stratify individuals by risk group and has promising applications in early diagnosis and clinical intervention [1, 2]. However, most PRS applications only leverage population-based cohorts and contrast each individual’s risk with that in the general population. It remains challenging to model and quantify within-family polygenic risk due to very limited samples in each family. Given parental data, it is statistically straightforward to model the Mendelian inheritance of any single genetic variant. However, PRS models include from dozens to millions of single-nucleotide polymorphisms (SNPs) that are scattered across the genome. These SNPs are often in linkage disequilibrium (LD) in the population. In addition, PRSs of family members are expected to have reduced variability compared to the population risk distribution due to limited crossover activities during meiosis. Such variability also varies across families. Therefore, modeling PRS within families requires careful considerations of phasing, linkage, and recombination.

Here, we introduce ORIGAMI (**O**ffspring **R**isk **I**nference through **GAM**ete s**I**mulation), a computational framework to simulate offspring genotypes using parental genetic data. ORIGAMI uses state-of-the-art, sex-specific genetic maps to simulate recombination events in phased parental genomes. Through extensive simulations and analyses of real data, we demonstrate that ORIGAMI simulations are computationally efficient and genetically realistic. By calculating PRS on simulated offspring for a couple, we can estimate the risk distribution for their potential children and compare this distribution to their observed children. To showcase the application of our approach, we incorporate ORIGAMI in polygenic transmission disequilibrium test (pTDT) [3] and demonstrate a substantial gain in statistical power after accounting for within-family PRS variability.

We applied ORIGAMI-informed pTDT to 7,805 trios of autism spectrum disorder (ASD) probands and healthy parents. ASD is an etiologically complex, clinically heterogeneous, and highly heritable neurodevelopmental disorder. Exome sequencing studies have identified more than 100 genes harboring ultra-rare or *de novo* variants for ASD [4]. Recently, heritability studies and GWAS implicated critical roles of common SNPs in the etiology of ASD [5, 6]. Additionally, multi-trait modeling revealed strong genetic correlations of ASD with various complex traits [7]. Compared to other approaches, pTDT leverages a robust, family-based design to assess the transmission disequilibrium of complex traits’ polygenic risk between healthy parents and ASD probands and shed important light on the shared genetics between ASD and other genetically correlated phenotypes [3]. In this paper, we showcased the improved power of pTDT after modeling within-family PRS variability and identified novel traits associated with the risk of ASD.

## Results

### Method overview

ORIGAMI takes parental VCF (or BCF) files as input and simulates offspring genotypes for the whole genome or a pre-specified list of SNPs (Figure 1). First, ORIGAMI simulates crossover locations in parental genomes. We used a chiasma interference model [8] implemented in Simcross (**URLs**) to simulate the chiasmata positions on each chromosome and then reduces the number of chiasmata positions to obtain the crossover locations. Between two neighboring crossover positions, we randomly select one phased haplotype to construct the gamete genome. Then, the offspring genome can be naturally constructed by combining the two gametes simulated from both parents. When needed, we repeat the procedure to generate a user-required number of offspring genomes in order to quantify the distribution of offspring polygenic risk. ORIGAMI outputs simulated genotypes into a set of PLINK files [9] which are readily analyzable in various downstream applications. We describe the detailed procedure in the **Methods** section.

**Figure 1.**
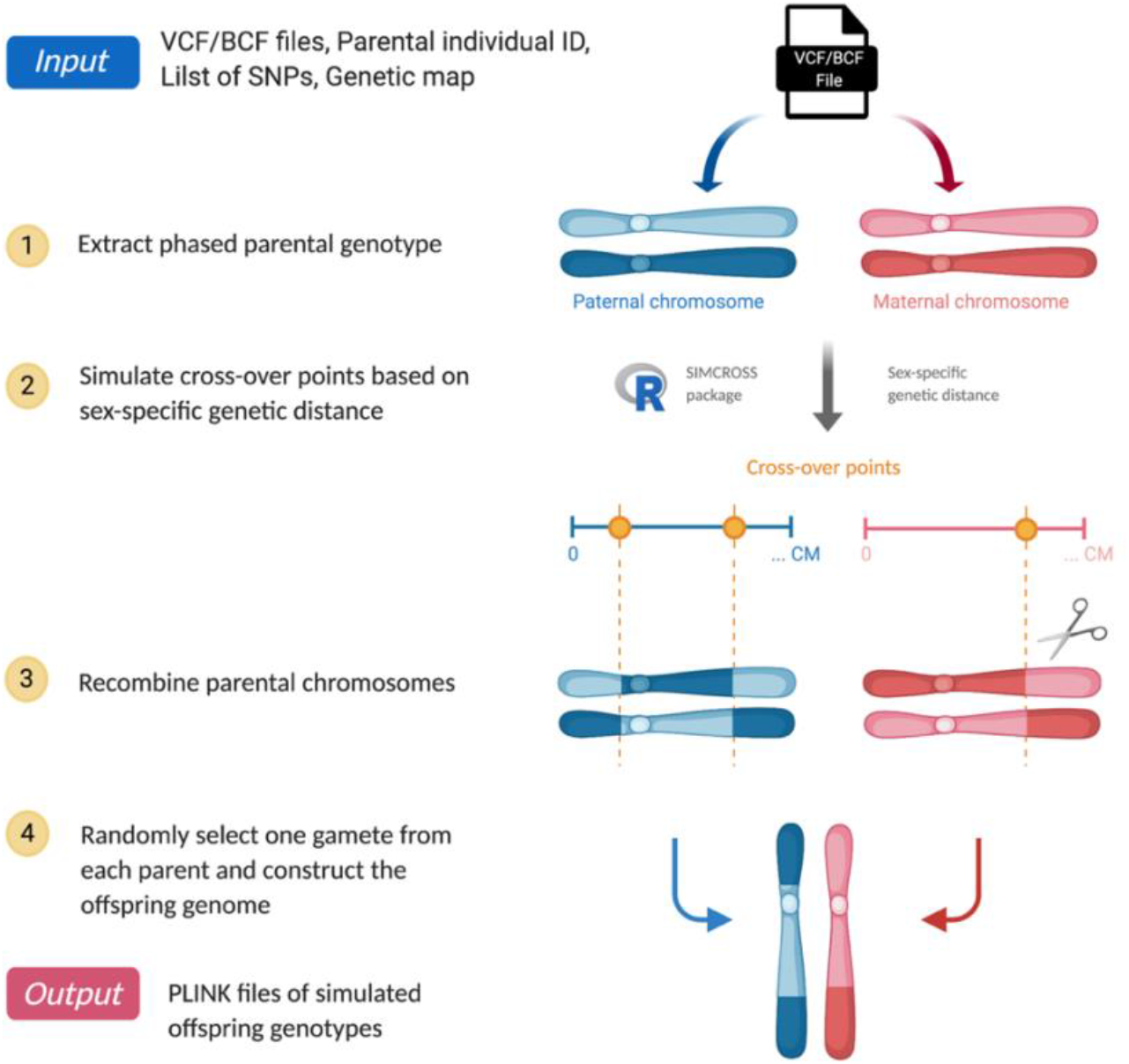
ORIGAMI workflow. Blue and red chromosomes represent paternal and maternal chromosomes, respectively.

We applied ORIGAMI to the spousal pairs in 7,805 ASD trios from three cohorts (**Methods**): the Autism Genome Project (AGP), the Simons Simplex Collection (SSC), and the Simons Foundation Powering Autism Research for Knowledge (SPARK). We simulated 100 pseudo children for each parental couple. We calculated PRS of ASD and other 67 complex traits (**Table S1**) for all parents and simulated offspring. Each PRS was statistically fine-tuned from GWAS summary statistics using PUMAS [10] (**Methods**) and standardized by cohort using the mean and standard deviation of parental scores.

### Validity of simulated offspring genotypes

We applied transmission disequilibrium test (TDT) to assess the transmission probability of each SNP between 15,610 parents and 780,500 simulated children. We performed TDT on three ASD cohorts separately and used METAL [11] to meta-analyze the associations. The results were consistent with the null distribution and suggested SNP-level transmission equilibrium from parents to their simulated offspring (Figure 2A).

**Figure 2.**
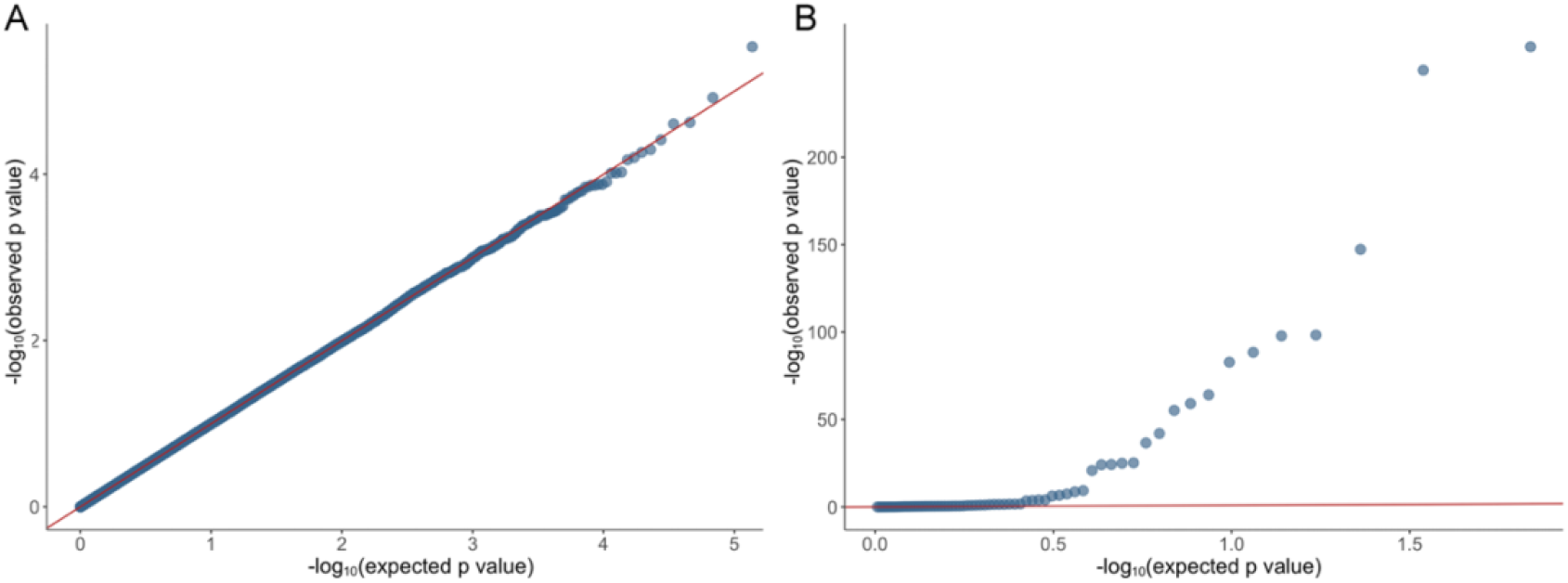
Evaluate the validity of simulated offspring genotype data. **(A)** Q-Q plot for TDT p-values. The x-axis shows the quantiles of the Unif(0,1) distribution. The y-axis shows the observed p-values in TDT. **(B)** We regressed child-parent PRS deviation on the estimated within-family PRS standard deviation. Q-Q plot for p-values is shown. The diagonal lines in both panels are highlighted.

Next, we compared PRS of 67 traits between simulated offspring and real samples in 3,242 trios with healthy parents and offspring (**Methods**). The parental average of PRS strongly associates with the average PRS of simulated offspring (**Table S2**), with 64 out of 67 traits showing correlations greater than 0.9. However, a unique and more important application of ORIGAMI is to quantify the variability of offspring PRS. We calculated the deviation of real siblings’ PRS from mid-parental PRS and associated these deviation values with the empirical standard deviation of simulated offspring’s PRS. The estimated PRS standard deviations were significantly associated with the absolute value of sibling-parent PRS deviation in 33 out of 67 traits (**Table S4**). The lower number of significant associations may be explained by the lack of real offspring samples per family, which poorly quantifies the variability of PRS within families. Still, these association p-values showed a significant deviation from the null (p=3.14e-12, Kolmogorov-Smirnov test; Figure 2B and **Table S3)**. Of note, we observed particularly strong associations in PRS with fewer SNPs (e.g., Alzheimer’s disease; **Figures S1a)**, but associations were also observed in more polygenic models (**Figure S1 b**). These results suggest that ORIGAMI effectively accounts for the linkage between SNPs and produces realistic genotype data for simulated offspring.

### Variability of genetic risk among siblings

Due to genetic similarity among family members, within-family variability of genetic risk is expected to be lower compared to the population-level variability. ORIGAMI-simulated offspring allow straightforward estimation of such variability. We calculated the PRS standard deviation among 100 simulated offspring of each family for 68 traits. We observed substantially reduced among-sibling PRS variability compared to that in the population (Figure 3A). The average among-sibling PRS standard deviation ranged from 46.6% to 80.8% of the standard deviation in the population across 68 traits (**Table S4)**. We also observed substantial heterogeneity of such variability across families. Using ASD risk score as an example, the among-sibling PRS standard deviation ranged between 42.2% and 91.8% (mean = 66.8%) of the population standard deviation in our data (Figure 3B). Across 68 traits, the range was 13.3% to 208.8% (**Table S3**). These results were consistent across multiple ASD cohorts (**Figure S2**) and also in simulated children for 2,715 spousal pairs in the Health and Retirement Study (HRS) (**Figure S3**). The between-family heterogeneity is significantly associated with the number of SNPs in PRS (p = 5.46e-06; Figure 3C). Traits with lower number of SNPs showed substantially flatter distributions of within-family standard deviation.

**Figure 3.**
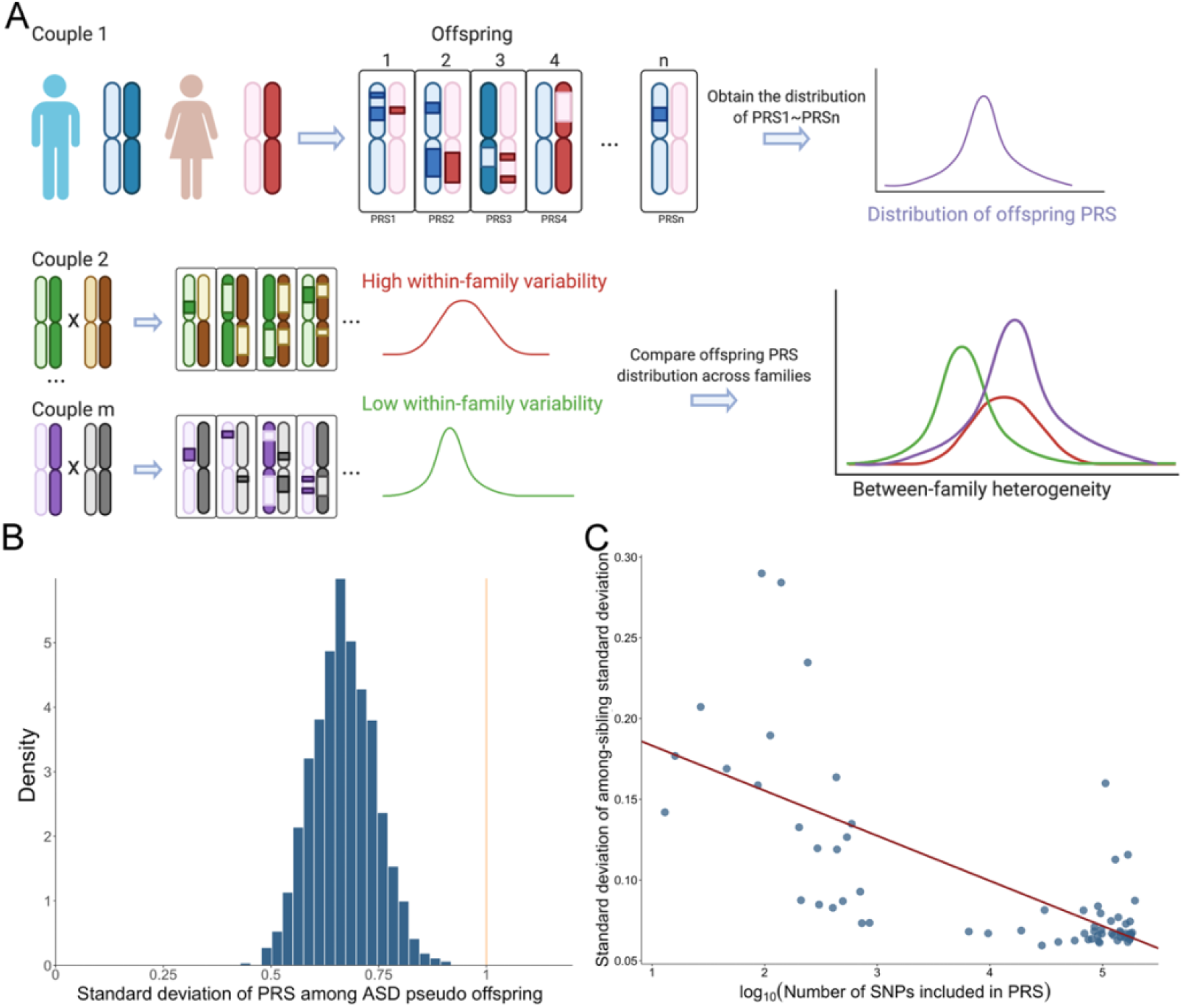
Within-family variability of polygenic risk. **(A)** a workflow for quantifying the variability of offspring PRS. We used ORIGAMI to generate n=100 pseudo offspring and estimated the offspring PRS distribution for each family. By comparing the PRS distribution across families, we quantified its between-family heterogeneity. **(B)** Histogram of the offspring PRS standard deviation. This distribution showcases the heterogeneous PRS variability across families. The orange line shows the standard deviation of PRS in the population. PRS was calculated using an ASD GWAS. **(C)** The heterogeneity of within-family PRS standard deviation is associated with number of SNPs in PRS. Each data point denotes a GWAS. Each value on the y-axis shows the standard deviation of a distribution similar to that in (B). The x-axis shows the log10-transformed number of SNPs. The fitted regression line is also shown.

### ORIGAMI-informed pTDT analysis

The standard pTDT analysis assumes an equal variance in the distribution of offspring PRS across different families (**Methods**). In practice, this assumption is most likely violated due to the variability in homozygosity at trait-informative loci, which can be affected by genetic drift, population stratification, and assortative mating [12]. Our results support that there exists substantial heterogeneity of within-family PRS variance (Figure 3B, **Figures S2-S3**). Instead of assuming an equal variance, we used ORIGAMI-simulated offspring to empirically estimate the offspring risk distribution, and standardized the pTDT test statistic (i.e., proband-parent PRS deviation) using within-family standard deviation estimates (Figure 4). This allows us to quantify the statistical evidence for polygenic transmission disequilibrium using a better-calibrated null distribution (**Methods**).

**Figure 4.**
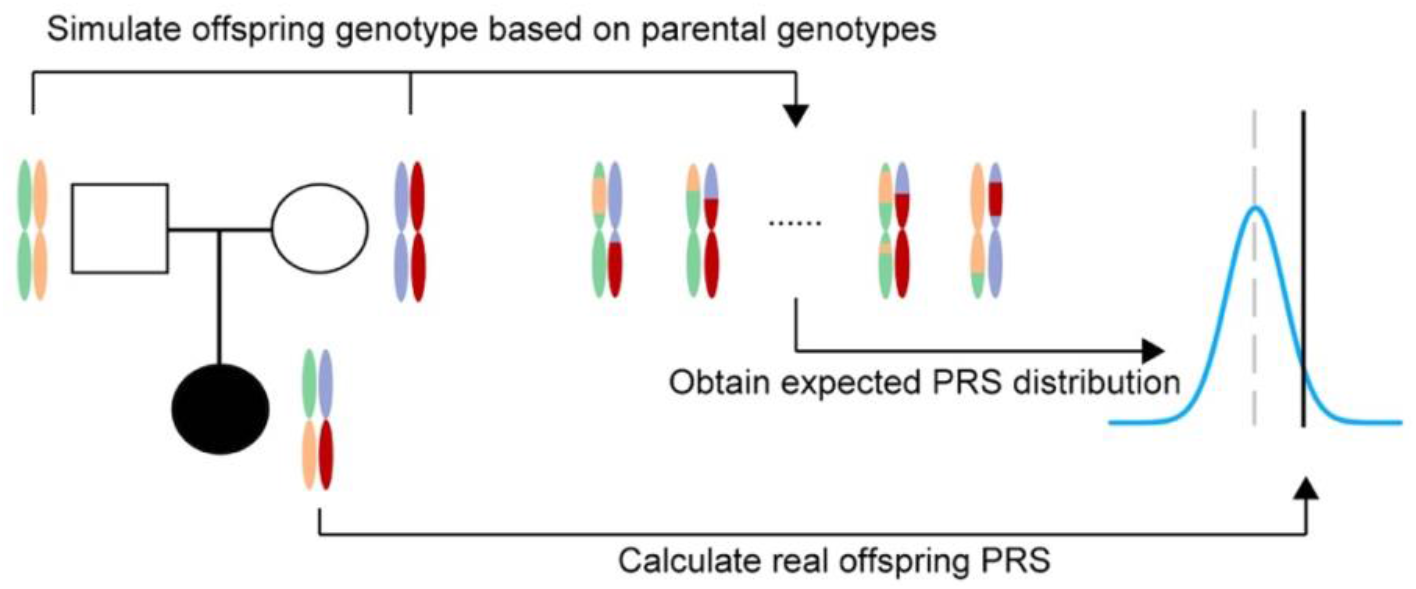
ORIGAMI-informed pTDT. We simulate offspring genotype data from parental genotypes and estimate the expected PRS distribution. pTDT is conducted by comparing the real offspring’s PRS with the expected PRS distribution.

We conducted simulations to compare the performance of standard pTDT and ORIGAMI-informed pTDT. We simulated PRS for 1000 trios under the assumption that the offspring PRS distribution could have distinct variance across families. We then empirically estimated the PRS standard deviation using 50, 100, 500, and 1000 simulated offspring per family and used the estimates as input in ORIGAMI-informed pTDT. Detailed simulation settings are described in the **Methods** section. Under transmission equilibrium, neither standard pTDT nor our approach showed inflated type-I error rates (Figure 5A). When PRS is associated with disease risk, our method showed substantially improved statistical power compared to the standard pTDT approach (Figure 5B). In addition, when sufficiently large (>50 in our analysis), the number of simulated offspring used to estimate within-family PRS variability did not influence type-I error or statistical power (**Figure S4**).

**Figure 5.**
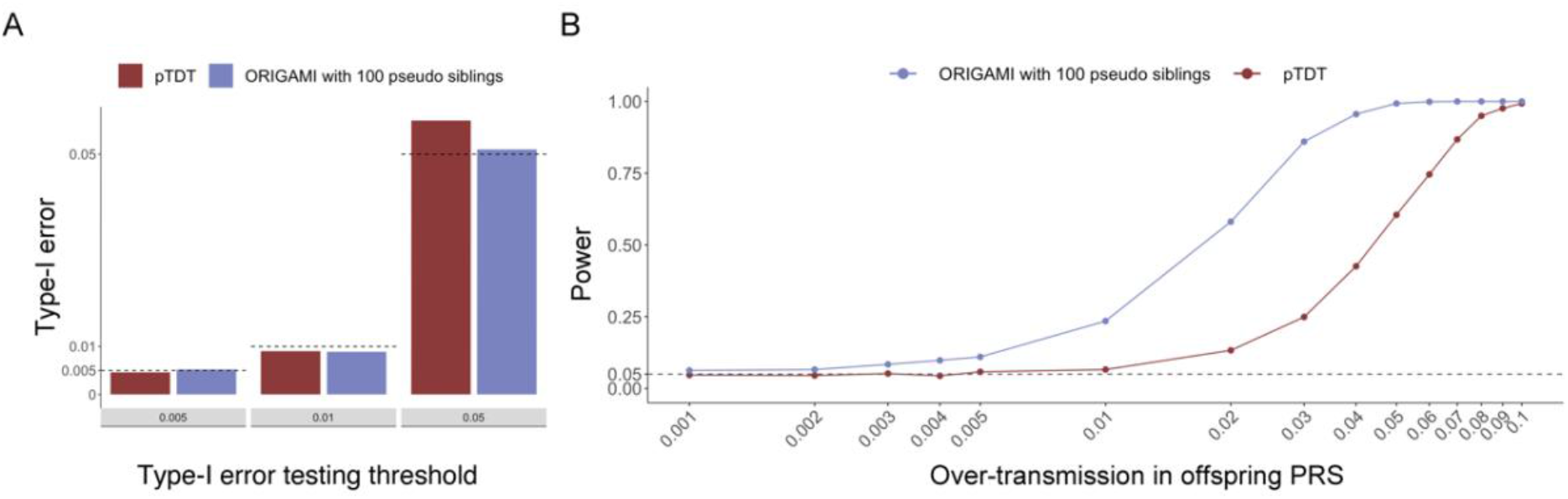
Statistical power and type-I error of standard and ORIGAMI-informed pTDT. **(A)** Type-I error of standard pTDT and ORIGAMI-informed pTDT with 100 simulated offspring. The dotted lines indicate significance thresholds. **(B)** Statistical power of standard pTDT and ORIGAMI-informed pTDT with 100 simulated offsprings. The scale of x-axis is log-transformed.

Next, we applied ORIGAMI-informed pTDT to assess the polygenic transmission equilibrium of ASD and 67 other complex diseases and traits in 7,805 ASD proband-parent trios. We identified 9 significant associations with false discovery rate (FDR) below 0.05 (Figure 6 and **Table S5**), 6 of which also reached statistical significance under Bonferroni correction (p<0.05/67=7.46e-4). We repeated the analysis in 3,242 healthy siblings-parent trios and no traits were significantly over- or under-transmitted (**Table S6**).

**Figure 6.**
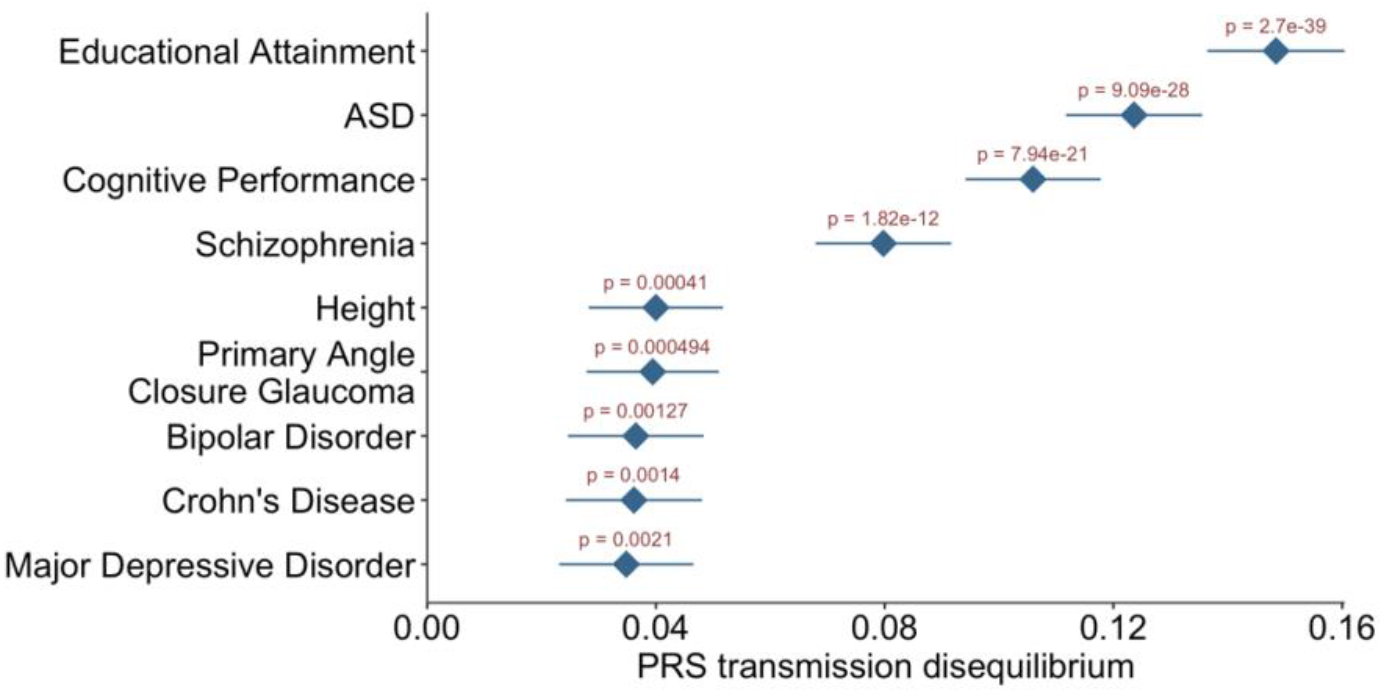
Complex traits showing significant transmission disequilibrium from healthy parents to ASD probands. Point estimates and standard errors were obtained from the meta-analysis of three ASD cohorts. X axis is the transmission between child and mid-parent PRS and Y axis shows the name of traits. Traits are order by p-value.

We confirmed the substantial over-transmission of ASD PRS from parents to probands with highly consistent effect sizes across three independent cohorts (p=9.09e-28; **Figure S5**; **Table S5**). Two additional traits that were previously identified to over-transmit in ASD probands [3], i.e., educational attainment (EA, p=2.70e-39) and schizophrenia (SCZ, p=1.82e-12), were also replicated in our analysis (**Figures S6-7**). In addition to the previously reported traits, we identified multiple significant novel associations including cognitive performance (p=7.94e-21), height (p=4.10e-4), primary angle closure glaucoma (p=4.94e-4), bipolar disorder (p=1.27e-3), and Crohn’s disease (p=1.40e-3) (**Figures S8-13**). All traits’ PRSs were over-transmitted from parents to probands. Notably, PRS of multiple neuropsychiatric traits, i.e., depressive symptoms (p=2.93e-3), attention deficit/hyperactivity disorder (ADHD; p=8.11e-3), and SCZ (p=0.034), were slightly under-transmitted to 3,242 healthy siblings (**Table S6**).

## Discussion

In this study, we introduced ORIGAMI, a computational framework to simulate genotypes of offspring from parental genomes. ORIGAMI uses sex-specific genetic maps to simulate crossover events in gametes and generates realistic offspring genotypes accounting for physical linkage. These features distinguish ORIGAMI from single-variant simulation approaches and make it particularly suitable for quantifying the polygenic risk within families. Comparing simulated offspring with the parents, we demonstrated transmission equilibrium at the SNP and PRS levels. We also demonstrated that the standard deviation of simulated PRS distribution is predictive for the real offspring-parents PRS deviation. Through estimating PRS variance among simulated offspring, ORIGAMI confirmed lower variability of polygenic risk within families and revealed substantial heterogeneity of such variability across families. Among 3,242 healthy trios and across 68 diseases and traits, the within-family PRS standard deviation varied from 13.3% to 208.8% of the PRS standard deviation in the population. The within-family PRS standard deviation is affected by multiple factors including the number of SNPs in PRS, weight values in PRS models, crossover frequency, and parental heterozygosity. ORIGAMI provides a computationally feasible approach to quantifying such variability.

The level of within-family risk variability directly affects the expected offspring-parent PRS deviation. Leveraging the empirical PRS distribution estimated from simulated offspring, we proposed an ORIGAMI-informed pTDT approach. Using simulations, we showed that our approach substantially improves the statistical power of pTDT without inflating the type-I error. We applied ORIGAMI-informed pTDT to a total of 7,805 trios of ASD probands and healthy parents from the SSC, AGP, and SPARK cohorts. We replicated previously reported over-transmission of ASD, SCZ, and EA PRS in ASD probands and identified significant over-transmission of PRS for 6 novel traits. No association reached statistical significance in 3,242 control trios which again demonstrated the well-calibrated type-I error of ORIGAMI-informed pTDT. We also note that pTDT results are generally consistent with genetic correlations between these traits and ASD [6, 13]. However, through leveraging family-based data, pTDT provides mechanistic insights into the shared genetics between ASD and correlated traits and produces results less susceptible to confounding or genetic associations mediated through family environment [14].

We also note that application of ORIGAMI is not limited to pTDT. As a novel tool that can prospectively quantify the polygenic risk of unborn offspring, ORIGAMI could have broad applications in risk stratification and genetic counseling. Theoretically, such information may also be used to help identify the ideal spouse who can maximize the probabilistic genetic prospect of future children. However, such applications are greatly complicated by ethnical concerns. In addition, our current understanding of complex traits’ genetic risk is insufficient to guarantee a causal effect on offspring phenotypes or even a reasonable level of predictive performance [15–17]. ORIGAMI also has some limitations. Our current implementation does not simulate sex chromosomes. Type of genetic variants simulated by ORIGAMI is limited to SNPs. Accuracy of ORIGAMI simulation also depends on phasing quality in the parental genomes. A future direction is to expand ORIGAMI to simulate diverse types of genetic variations, including variants on sex chromosomes, and to account for uncertainty in phasing.

Taken together, ORIGAMI is a novel, powerful, and computationally efficient framework for offspring genotype simulation. We showcased its application in pTDT inference and demonstrated substantially improved statistical power after accounting for within-family risk variability. The unique features of quantifying the offspring genetic risk at the genome-wide level makes ORIGAMI an appealing tool in human genetics research and prospective risk inference.

## Methods

### Sample information

Data from the AGP cohort were accessed through dbGaP (accession: phs000267) which consisted of 7,880 samples, including 2,188 trios of ASD probands and healthy parents. The SSC and SPARK cohorts were accessed through the Simons Foundation Autism Research Initiative (SFARI). Details on these data can be found on the SFARI website (URLs) and have been previously described [6, 18, 19]. We removed samples with a genotype missing rate > 0.05 and removed SNPs that have a genotype call rate < 0.95, significant deviation from Hardy-Weinberg equilibrium (p<1e-06), or a minor allele frequency < 0.01. We also removed overlapping individuals across cohorts using genetic relationship coefficients estimated by GCTA [20]. We only included individuals with self-reported European ancestry in the analyses. After the quality control (QC) and filtering process, 2,188, 1,794 and 3,823 proband trios remained in the AGP, SSC, and SPARK cohorts, respectively. 1,432 and 1,813 control trios of healthy siblings and parents remained in the SSC and SPARK cohorts. We used the UCSC liftover software (**URLs**) to lift over the genome build to hg19. We used the Michigan Imputation server [21] to phase and impute all genotype data to the Haplotype Reference Consortium reference (version r1.1 2016). SNPs with imputation quality score < 0.8 and minor allele frequency < 0.01 were removed from the analysis. After post-imputation QC, 7,260,224 SNPs remained in the AGP. The 1Mv1, 1Mv3, and Omni2.5 cohorts (defined by genotyping arrays) in the SSC had 7,298,961 SNPs, 7,029,817 SNPs, and 6,866,248 SNPs, respectively. 7,031,717 SNPs remained in the SPARK cohort.

To validate results in a population cohort, we also simulated 100 children for each spousal couple (N=2,715 couples) in the HRS. The HRS is a nationally representative, longitudinal panel study of individuals over the age of 50 and their spouses conducted by the University of Michigan [22]. Imputed HRS genotype data were accessed through dbGap (phs000428). We calculated PRS of ASD in HRS using the same GWAS summary statistics used in three ASD cohorts.

### Statistical and computational details of the ORIGAMI framework

ORIGAMI is a Linux-based software tool. Core functions of ORIGAMI were implemented in R. ORIGAMI takes phased VCF or BCF files as the input. It first uses bcftools [23] to extract autosomal genotypes from paternal and maternal files and then simulates crossover locations in the parental genomes. For each chromosome, we used a chi-square chiasma interference model [8] which accounts for genetic distance and interference. We set the interference number to be 3 and used sex-specific genetic maps [24] to simulate chiasma counts in each parent:

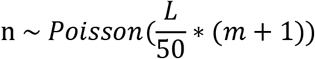

where n indicates the number of chiasmata, m is the interference number, L is the maximum genetic distance in the chromosome. After sampling chiasma counts, we simulate the first chiasma position indicator *n_f_*, which is a random number taking values from 0 to m. *n_f_* = 2 means the first chiasma is the second simulated chiasma position, so the first simulated chiasma position is discarded. f stands for first here. Then the final number of chiasma points is *n* – *n_f_* + 1. If *n_f_* ≥ *n* + 1, then there will be no interference. Otherwise, we simulate *n – n_f_* + 1 chiasma position. Here, we assume that the chiasma position follows a uniform distribution *Unif* (0, *L*). Here, an interference number of 3 means that there are about 52 chiasmata for males and 80 for females in the whole genome on average. Each simulated chiasma position will be deleted with a probability of 50% since tetrads enjoying one or more crossovers in a marked interval only produce 50% recombinant haploid products when lacking chromatid interference [8]. Then, crossover events will happen at the remaining chiasma positions.

For each autosome in a parental genome, ORIGAMI recombines phased haplotypes at the crossover positions and generates a pair of recombined chromosomes, from which we randomly select one to be the gamete chromosome. Finally, we create a pseudo offspring genome by combining two simulated gametes from both parents. Repeating the procedure, ORIGAMI generates a user-specified number of offspring genomes for each spousal pair, and outputs a set of binary PLINK [9] files for merged offspring genotypes. The current ORIGAMI implementation does not simulate sex chromosomes.

### GWAS summary statistics and PRS calculation

We obtained GWAS summary statistics for 68 complex traits from the literature and public repositories. Details on these studies are summarized in **Table S1.** We computed PRSs in the AGP, SSC, and SPARK cohorts. SNPs with strong LD were clumped in PLINK [9]. We used the genotype data of European descendants in 1000 Genome Project Phase III cohort [25] as the LD reference and clumped summary-level data by an LD window size of 1Mb and pairwise *r*^2^ thresholds of 0.1. In addition, we fine-tuned the optimal p-value cutoff for each PRS through summary statistics-based cross-validation implemented in PUMAS [10]. The optimal p-value cutoffs and maximal predictive *R*^2^ are reported in **Table S1**. All PRS computations were carried out using PRSice-2 under the PUMAS-tuned p-value thresholds [26].

### Validating simulated offspring genotype data

We performed TDT analysis for 137,021 SNPs included in all three ASD cohorts and the ASD PRS model. We performed TDT in the AGP, SSC and SPARK cohorts separately and meta-analyzed the associations using METAL [11]. We also tested the association between the standard deviation of within-family PRS standard deviation (i.e., between-family heterogeneity) and the log10-transformed number of SNPs included in the PRS models. The number of SNPs in a PRS will be affected by the p value cutoff fine-tuned by PUMAS.

### ORIGAMI-informed pTDT

In the standard pTDT approach [3], PRS deviation in the *i^th^* trio is calculated as

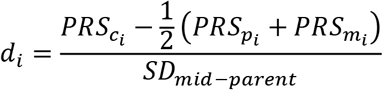

Here, *PRS_c_i__, PRS_p_i__*, and *PRS_m_i__* denote the child, paternal, and maternal PRSs in the *i^th^* trio, respectively. *SD_mid–parent_* denotes the standard deviation of mid-parental PRS values (i.e., (*PRS_p_i__* + *PRS_m_i__*)/2) across trios. This approach implicitly assumes the variance of offspring PRS to be same across families, thus dividing the same standard deviation estimate to standardize the test statistic. To evaluate the difference between pTDT deviation and 0, the pTDT test statistics is calculated as

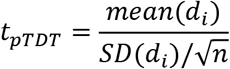

Under this assumption, the pTDT test statistics follows a t distribution under the null hypothesis (i.e., transmission equilibrium of PRS). However, we demonstrated that the variance of offspring PRS could vary greatly in different families, which violates the assumption in t test and reduces statistical power of pTDT.

The ORIGAMI-informed pTDT replaces the denominator with the estimated standard deviation of offspring PRS in each family:

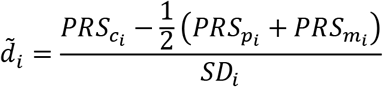

where *SD_i_* denotes the standard deviation of offspring PRS of the *i^th^* family. Importantly, we allow different families to have distinct variance values in the offspring PRS distribution. The empirical standard deviation of simulated children’s PRS in the *i^th^* family provides an estimate of *SD_i_*. Under this relaxed assumption, we can calculate the ORIGAMI-informed pTDT test statistics as

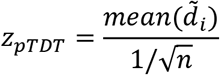

Under the null, the ORIGAMI-informed pTDT test statistic follows a standard normal distribution asymptotically.

### Simulation settings

We simulated PRS data from 1000 families. We randomly sampled 1000 data points from a normal distribution as the standard deviation of PRS these families.

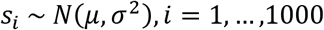

Here we set *μ* = 0.65 and *σ* = 0.3 based on the among-sibling variability of 67 PRSs in our analyses. Parental PRSs were randomly sampled from the standard normal distribution *N*(0,1). For type-I error evaluation, we generated 50, 100, 500, and 1000 “healthy” offspring PRS values for each family under transmission equilibrium from 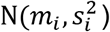, where *m_i_* is the average of parental PRS in the i^th^ family. To assess statistical power, we generated 50, 100, 500, and 1000 “proband” PRS values for each family under transmission disequilibrium. More specifically, we sampled offspring PRSs from 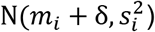 for the i^th^ family. Here, *s_i_* still follows *N*(0.65, (0.3)^2^) as described above. δ is the effect size of transmission disequilibrium which ranged from 10^-5^ to 0.1. We empirically estimated the standard deviation *s_i_* for each family using simulated offspring data and calculated pTDT deviation. We used t distribution and standard normal distribution as the null for standard and ORIGAMI-informed pTDT, respectively. We repeated the process 1000 times and quantified type-I error rate and statistical power as the proportion of repeats that reached statistical significance at a given threshold.

## URLs

AGP (https://www.ncbi.nlm.nih.gov/projects/gap/cgi-bin/study.cgi?study_id=phs000267.v5.p2);

SSC (https://www.sfari.org/resource/simons-simplex-collection/);

SPARK (https://www.sfari.org/resource/spark/);

PLINK (https://zzz.bwh.harvard.edu/plink/)

PUMAS (https://github.com/qlu-lab/PUMAS)

Simcross (https://github.com/kbroman/simcross)

PRSice-2 (https://www.prsice.info/)

Liftover (https://genome.ucsc.edu/cgi-bin/hgLiftOver)

## Code availability

The ORIGAMI software is publicly available at https://github.com/qlu-lab/ORIGAMI

## Author contribution

Q.L. conceived and designed the study.

J.C. and Q.L. developed the statistical framework.

J.C. and J.Y. implemented the software.

J.C. performed the statistical analysis.

Y.W., J.Y. and K.H. conducted data processing and quality control.

Z.Z. and J.C. performed the PUMAS p-value cut off optimization and calculated the PRS.

Z.N. assisted the polygenic transmission disequilibrium analysis.

Z.Z. and Z.N. clumped the GWAS datasets.

Q.L. and J.F. advised on statistical and genetic issues.

J.C. and Q.L. wrote the manuscript.

All authors contributed in manuscript editing and approved the manuscript.

## Acknowledgements

We are grateful to all the families participating in the Autism Genome Project (AGP), the Simons Simplex Collection (SSC), and the Simons Foundation Powering Autism Research for Knowledge (SPARK) study. This project was supported by the Clinical and Translational Science Award (CTSA) program, through the NIH National Center for Advancing Translational Sciences (NCATS), grant UL1TR000427. We gratefully acknowledge use of the facilities of the Center for Demography of Health and Aging at the University of Wisconsin-Madison, funded by NIA Center Grant P30 AG017266. We also acknowledge research support from the University of Wisconsin-Madison Office of the Chancellor and the Vice Chancellor for Research and Graduate Education with funding from the Wisconsin Alumni Research Foundation and the Waisman Center pilot grant program at the University of Wisconsin-Madison. This research uses data from HRS which is supported by National Institute on Aging Grants U01AG009740, RC2AG036495, and RC4AG039029 and is conducted by the University of Michigan. We thank Drs. Jakob Grove and Elise Robinson for sharing the GWAS summary statistics based on the iPSYCH cohort. We thank members of the Social Genomics Working Group at University of Wisconsin for helpful comments. We thank Dr. Karl Broman for helpful discussions on the simulation of crossover events. The workflow figures were created using Biorender.

