## Supplementary figure for "Gamete simulation improves polygenic transmission disequilibrium analysis"

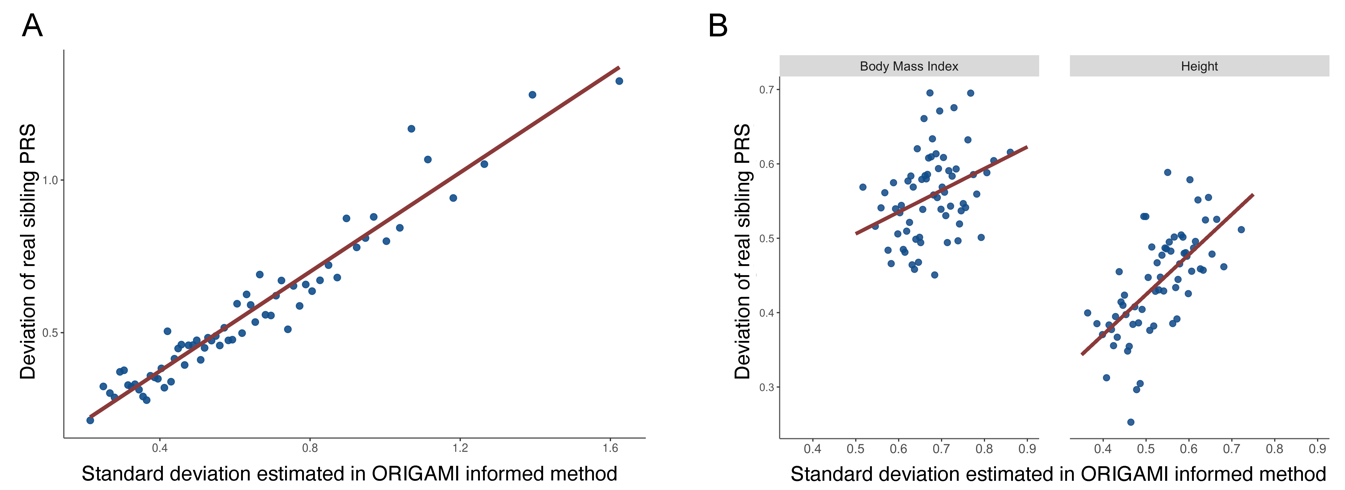

**Supplementary Figure 1. Within-family PRS standard deviation is associated with the real child-parent PRS deviation.** Parental PRS, real offspring PRS, and simulated offspring PRS were all standardized by the mean and standard deviation of parental PRS. The figure shows binned scatterplots in which each plotted point reflects the average x and y coordinates for a bin of 50 individuals. The red regression lines were fitted on raw data. The x-axis shows the within-family PRS standard deviation calculated from 100 pseudo offspring for each family. **(A)** Association results for Alzheimer’s disease PRS. **(B)** Association results for BMI and height PRS.

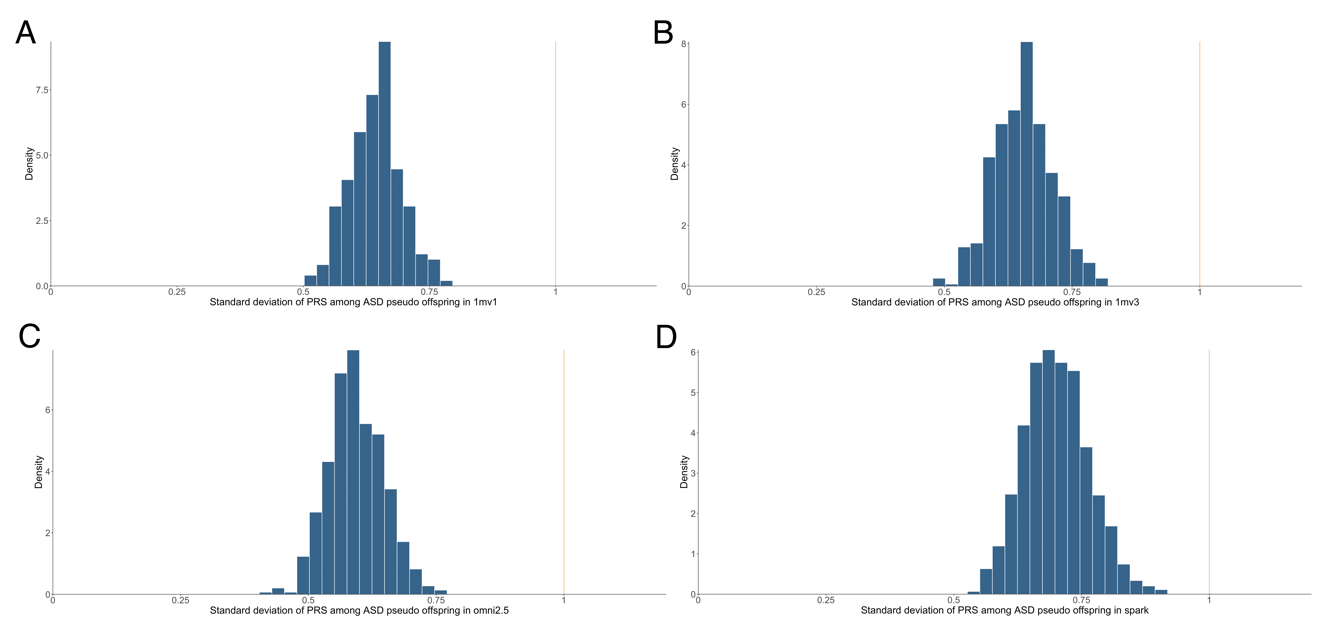

**Supplementary Figure 2. Histogram of the offspring PRS standard deviation across ASD cohorts**. This distribution showcases the heterogeneous PRS variability across families in (A) SSC 1mv1, (B) SSC 1mv3, (c) SSC omni2.5, and (D) SPARK cohorts. The pattern of offspring PRS standard deviation remained similar across different cohorts. The orange line shows the standard deviation of PRS in the population. PRS was calculated using the same ASD GWAS used for Figure 3B.

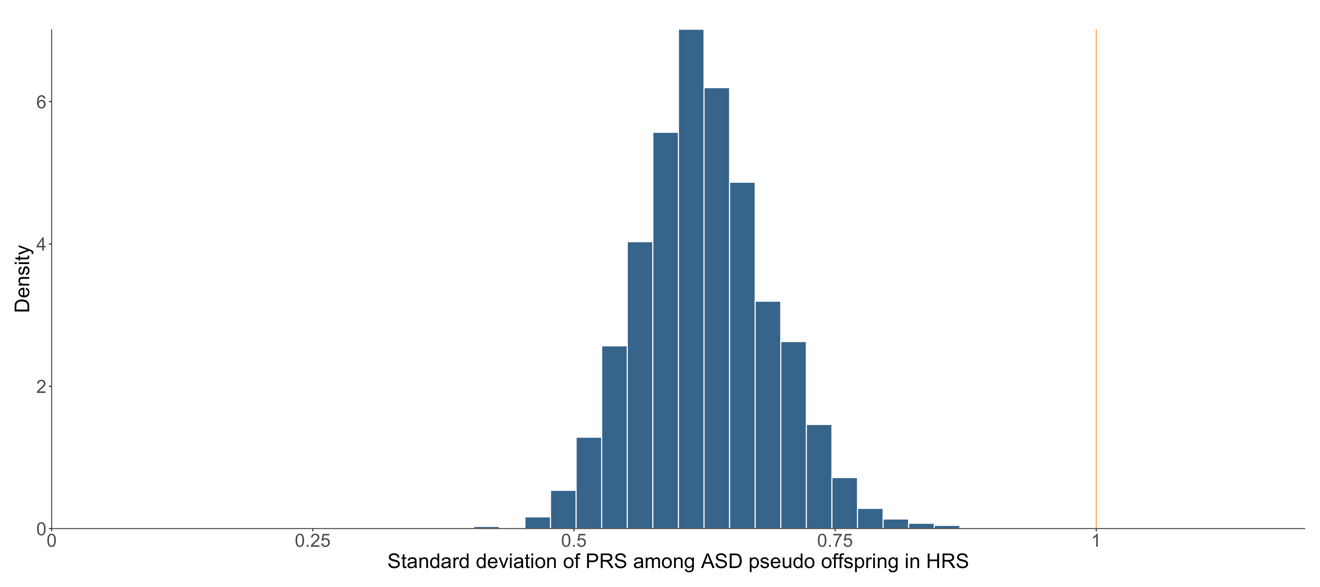
**Supplementary Figure 3. Histogram of the offspring PRS standard deviation in HRS.** This distribution showcases the heterogeneous PRS variability across families in the HRS cohort. The pattern of offspring PRS standard deviation is similar compared to the ASD cohorts. The orange line shows the standard deviation of PRS in the population. PRS was calculated using the same ASD GWAS used for Figure 3B.

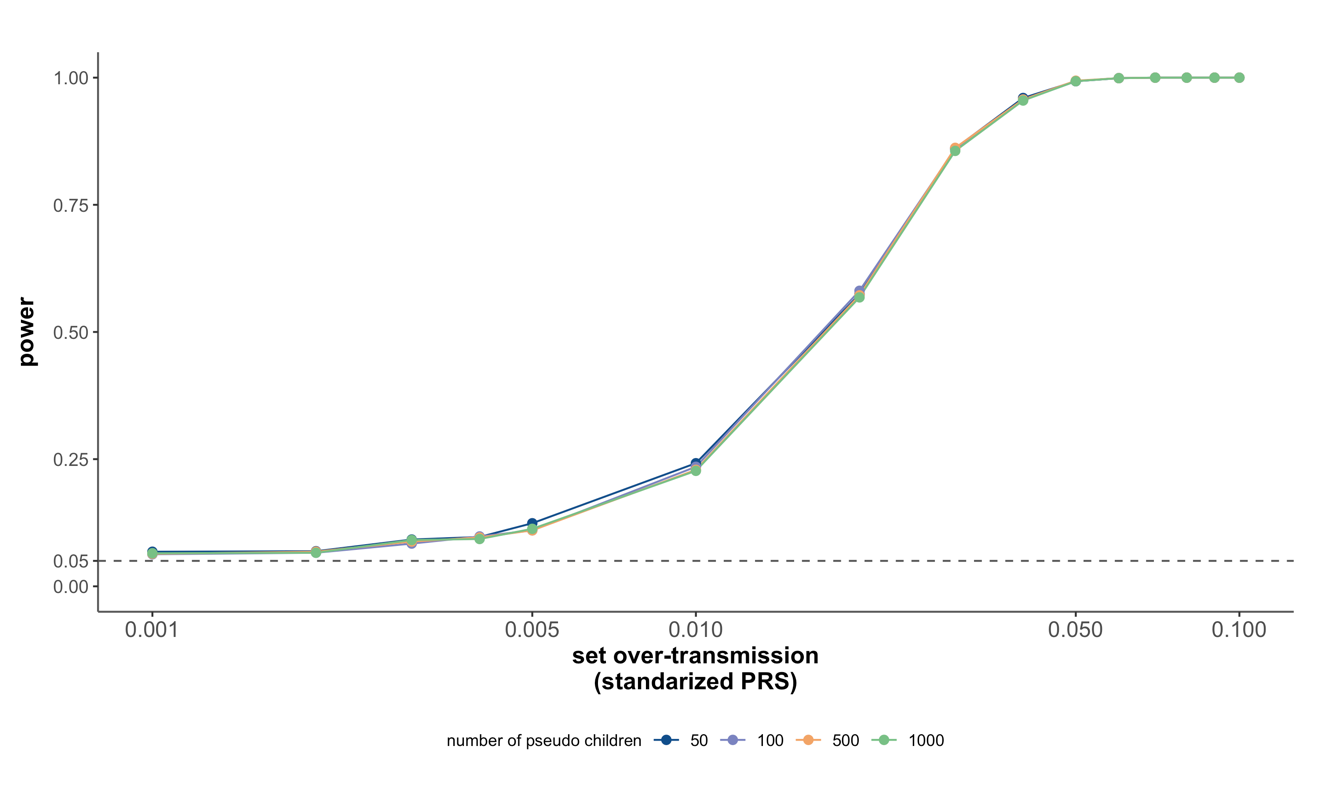

**Supplementary Figure 4. Statistical power of ORIGAMI-informed pTDT analysis.** We compared the power of ORIGAMI-informed pTDT analysis using different numbers of pseudo offspring in the simulation. The results were robust to the number of offspring.

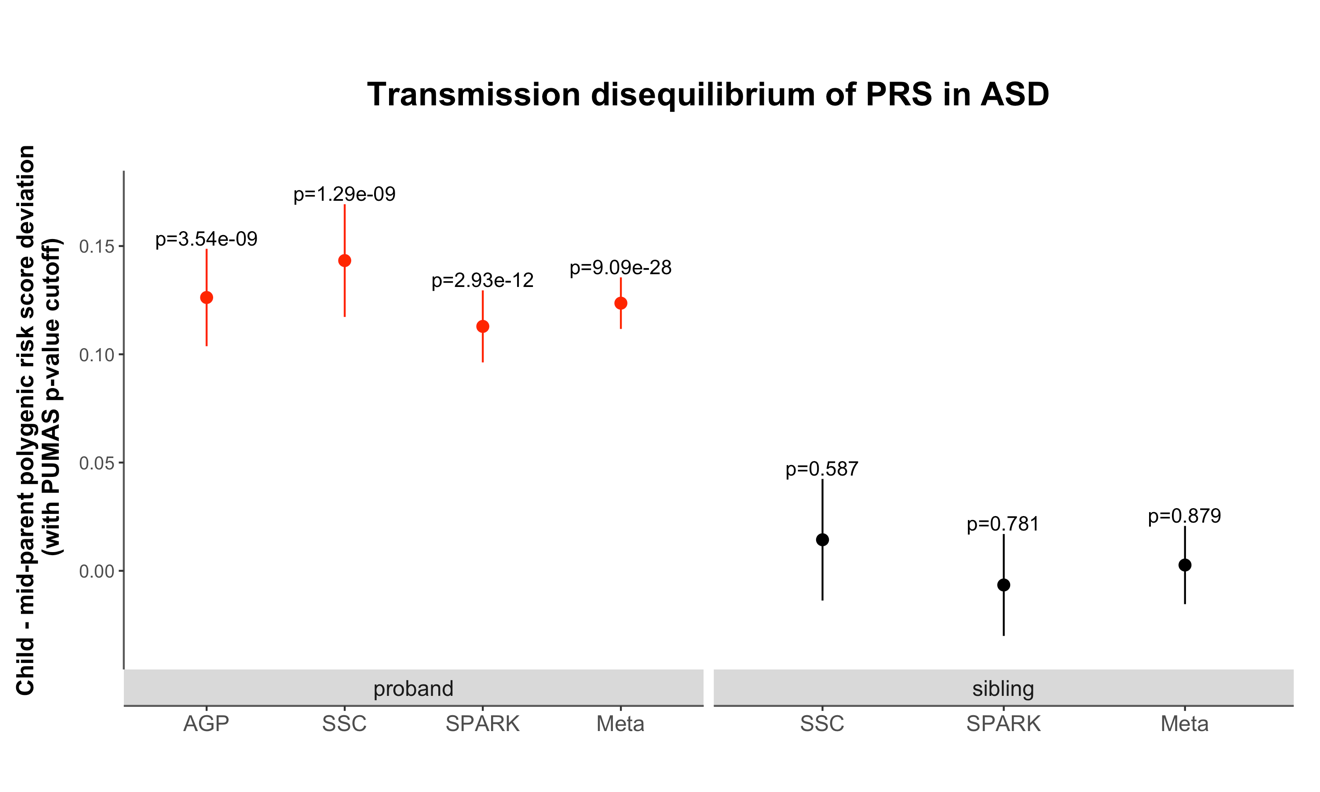
**Supplementary Figure 5. Transmission disequilibrium of PRS in different cohorts in ASD using ORIGAMI informed pTDT analysis.** Transmission disequilibrium was quantified by the ORIGAMI informed pTDT analysis with PUMAS p-value cut off. Results in probands and unaffected siblings are highlighted in different colors. The mean pTDT deviation and the SE are shown. P-values are labeled above each interval.

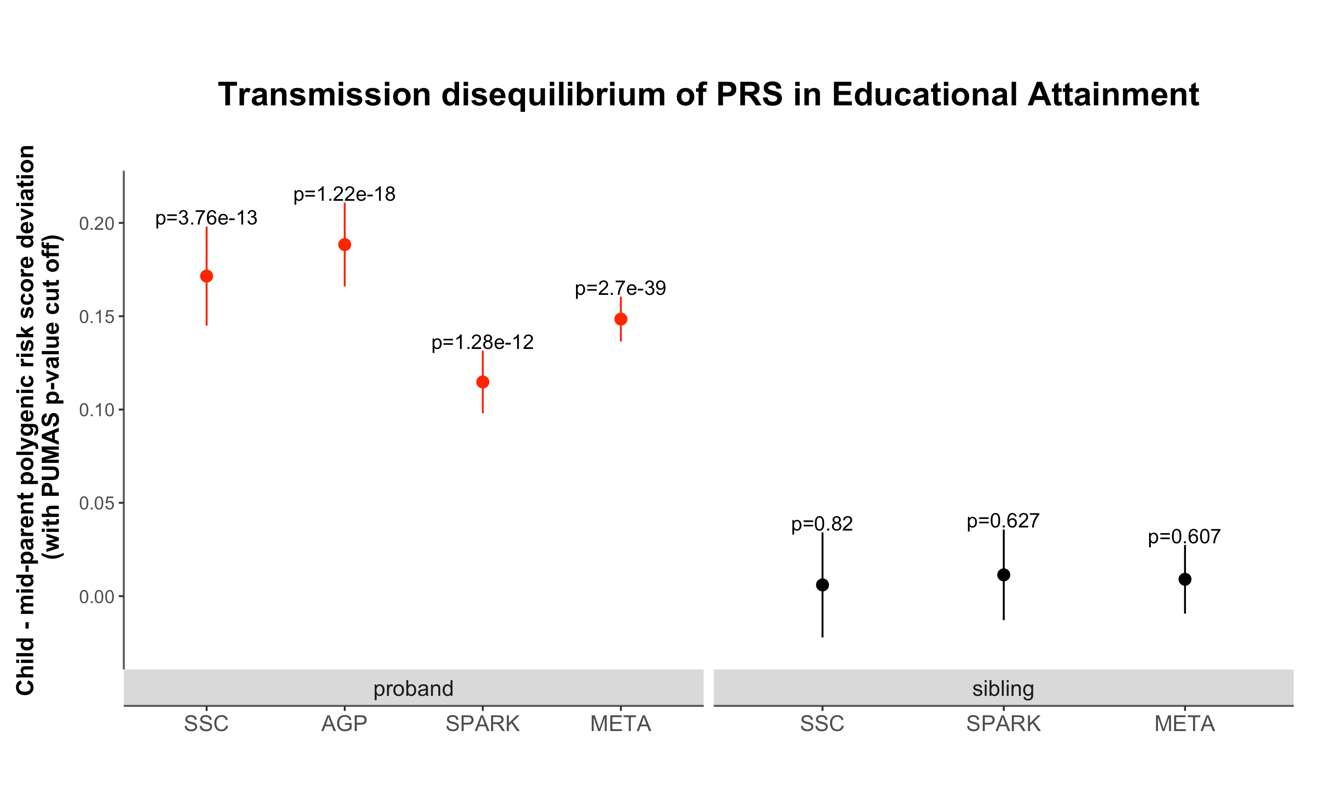

**Supplementary Figure 6. Transmission disequilibrium of PRS in different cohorts in educational attainment using ORIGAMI informed pTDT analysis.** Transmission disequilibrium was quantified by the ORIGAMI informed pTDT analysis with PUMAS p-value cut off. Results in probands and unaffected siblings are highlighted in different colors. The mean pTDT deviation and the SE are shown. P-values are labeled above each interval.

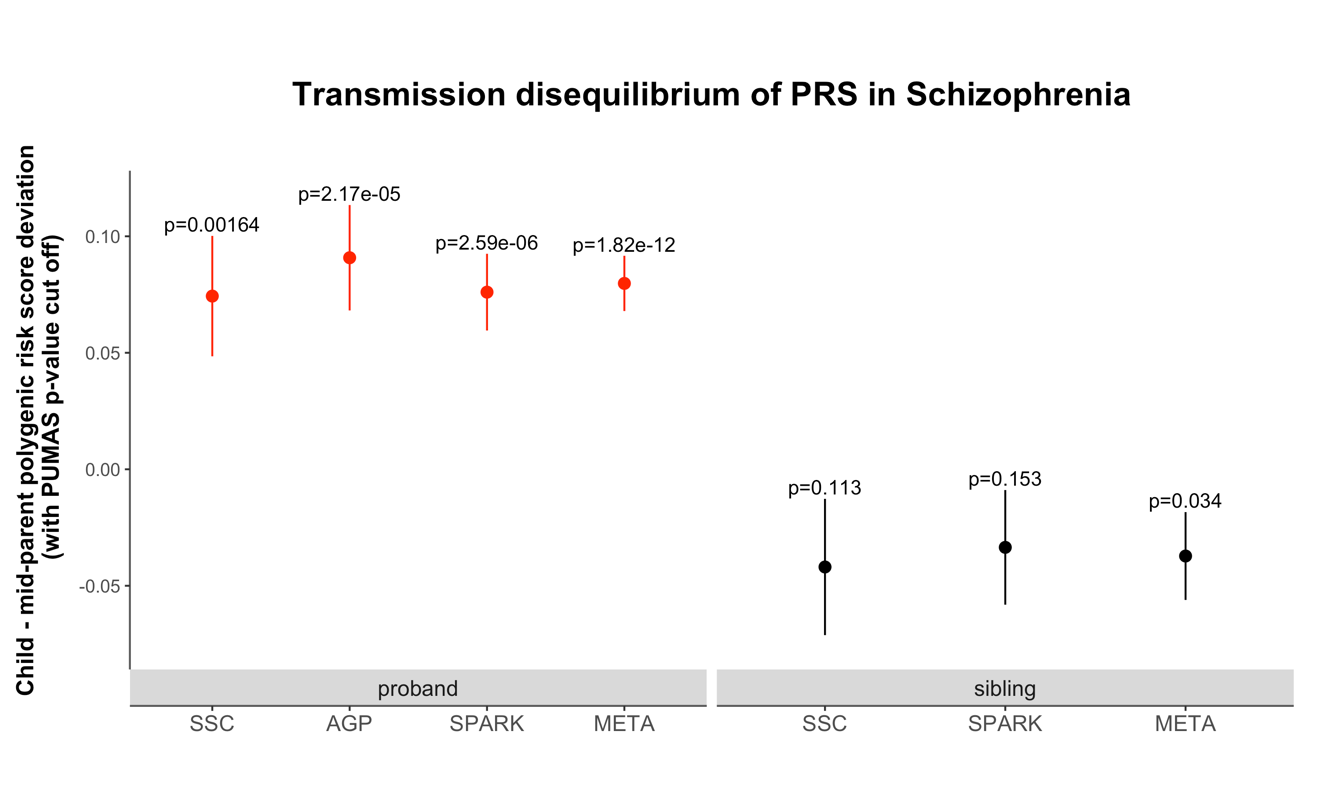

**Supplementary Figure 7. Transmission disequilibrium of PRS in different cohorts in schizophrenia using ORIGAMI informed pTDT analysis.** Transmission disequilibrium was quantified by the ORIGAMI informed pTDT analysis with PUMAS p-value cut off. Results in probands and unaffected siblings are highlighted in different colors. The mean pTDT deviation and the SE are shown. P-values are labeled above each interval.

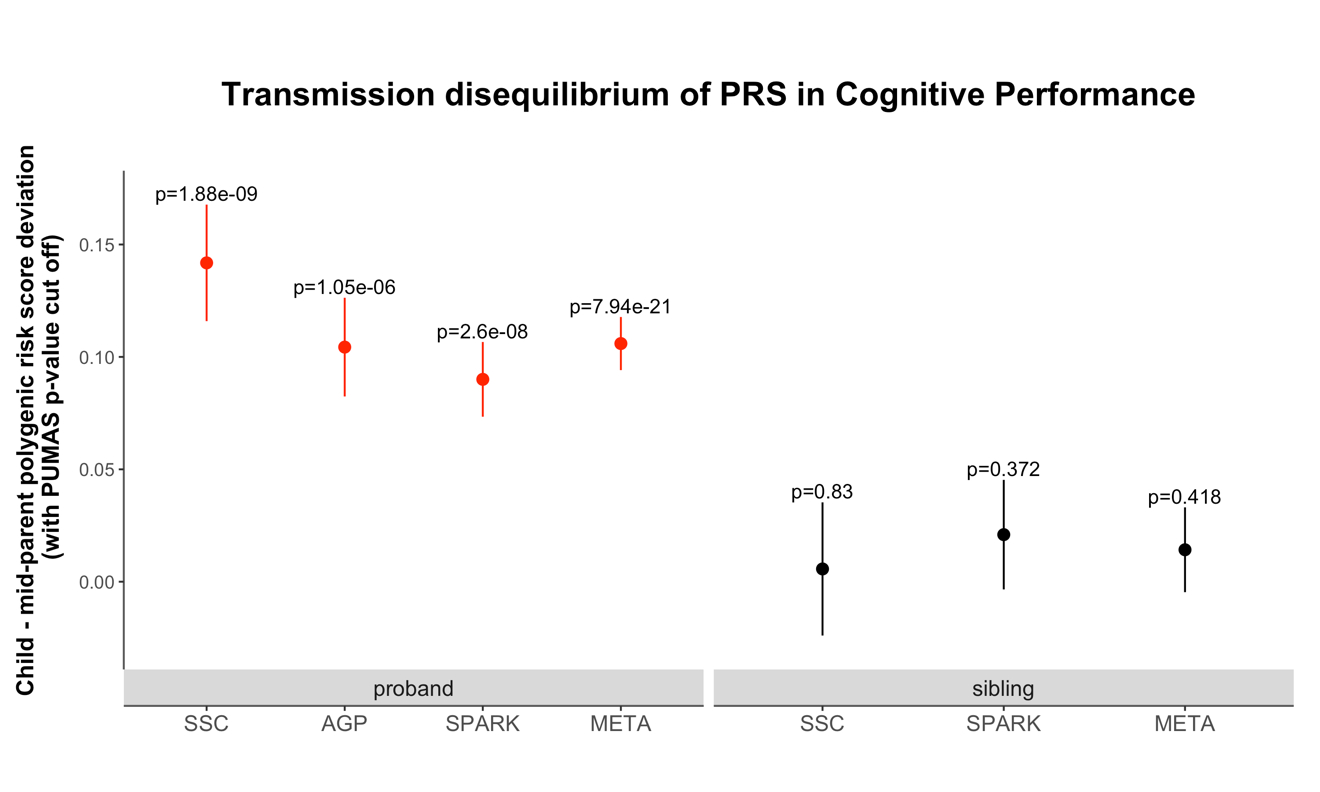

**Supplementary Figure 8. Transmission disequilibrium of PRS in different cohorts in cognitive performance using ORIGAMI informed pTDT analysis.** Transmission disequilibrium was quantified by the ORIGAMI informed pTDT analysis with PUMAS p-value cut off. Results in probands and unaffected siblings are highlighted in different colors. The mean pTDT deviation and the SE are shown. P-values are labeled above each interval.

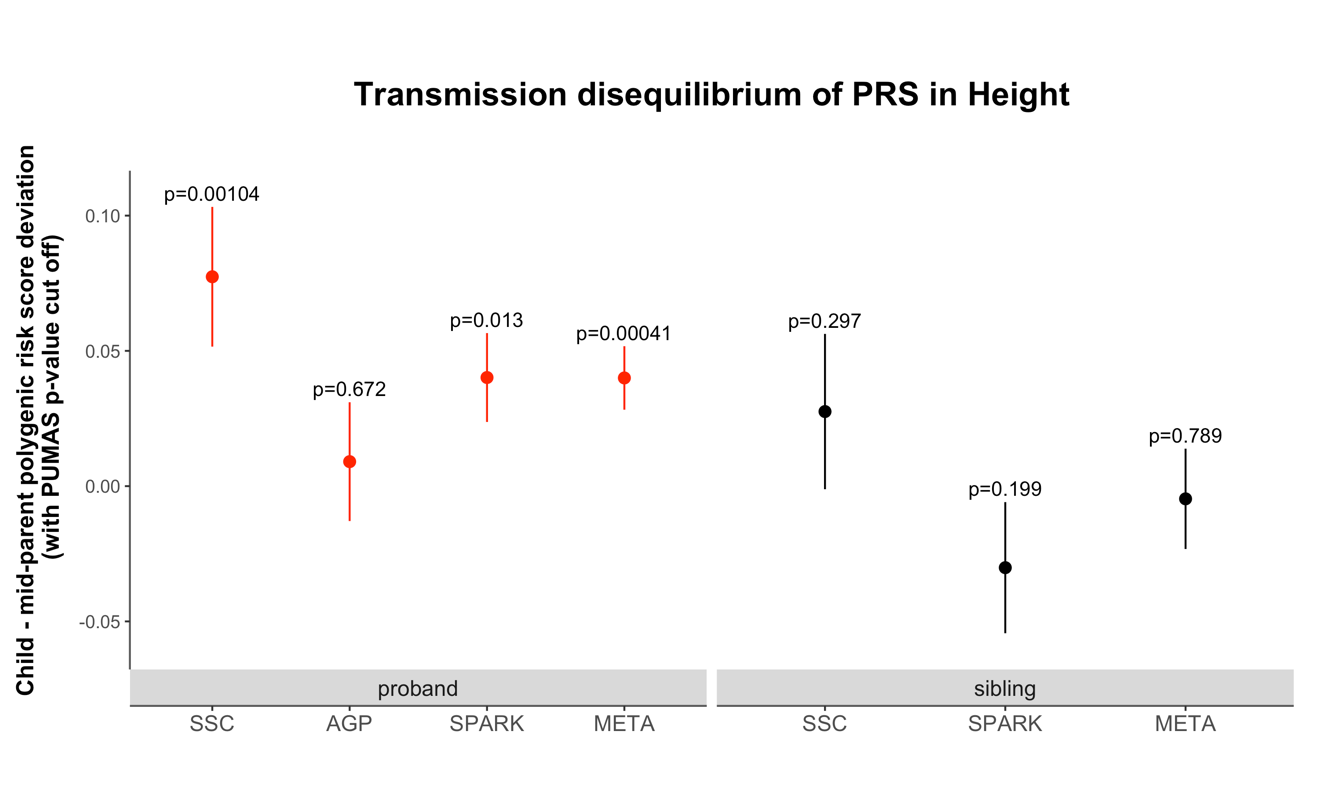

**Supplementary Figure 9. Transmission disequilibrium of PRS in different cohorts in height using ORIGAMI informed pTDT analysis.** Transmission disequilibrium was quantified by the ORIGAMI informed pTDT analysis with PUMAS p-value cut off. Results in probands and unaffected siblings are highlighted in different colors. The mean pTDT deviation and the SE are shown. P-values are labeled above each interval.

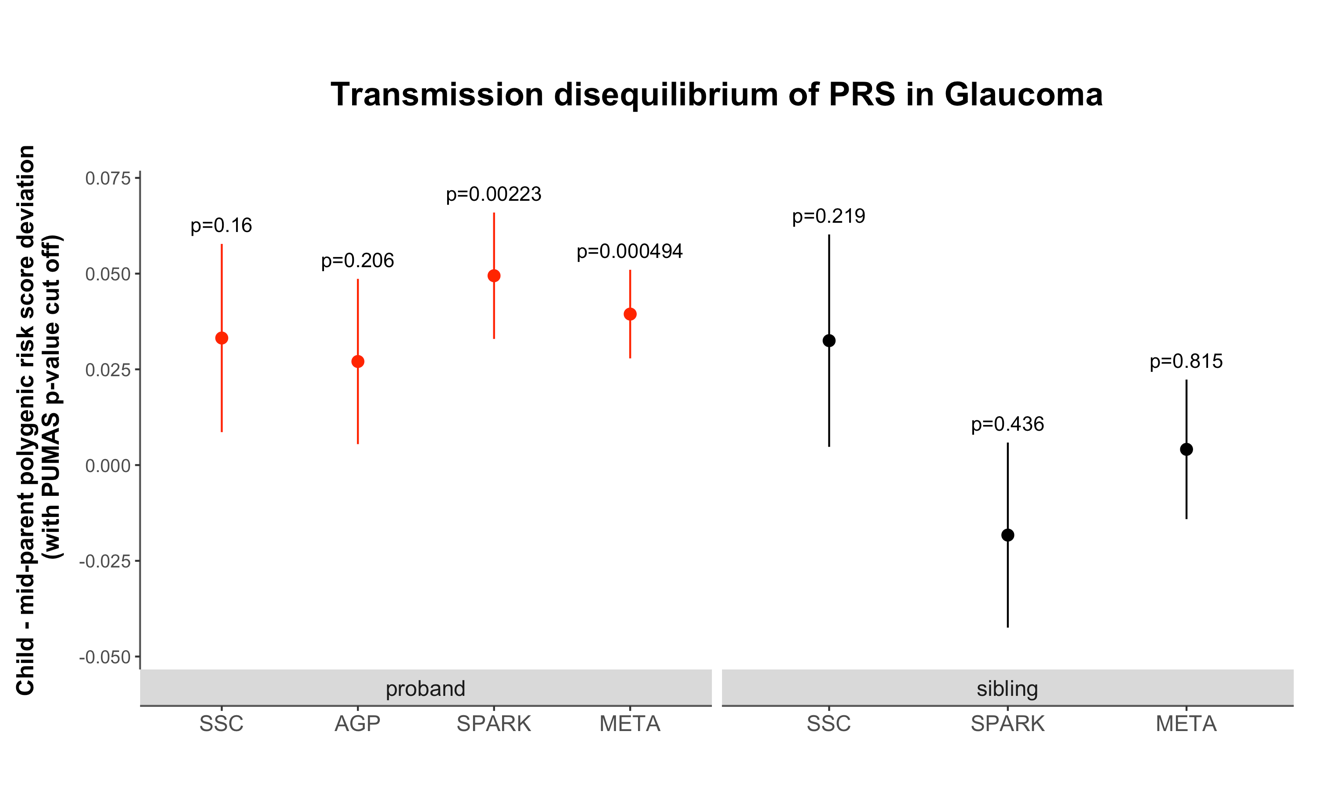

**Supplementary Figure 10. Transmission disequilibrium of PRS in different cohorts in glaucoma using ORIGAMI informed pTDT analysis.** Transmission disequilibrium was quantified by the ORIGAMI informed pTDT analysis with PUMAS p-value cut off. Results in probands and unaffected siblings are highlighted in different colors. The mean pTDT deviation and the SE are shown. P-values are labeled above each interval.

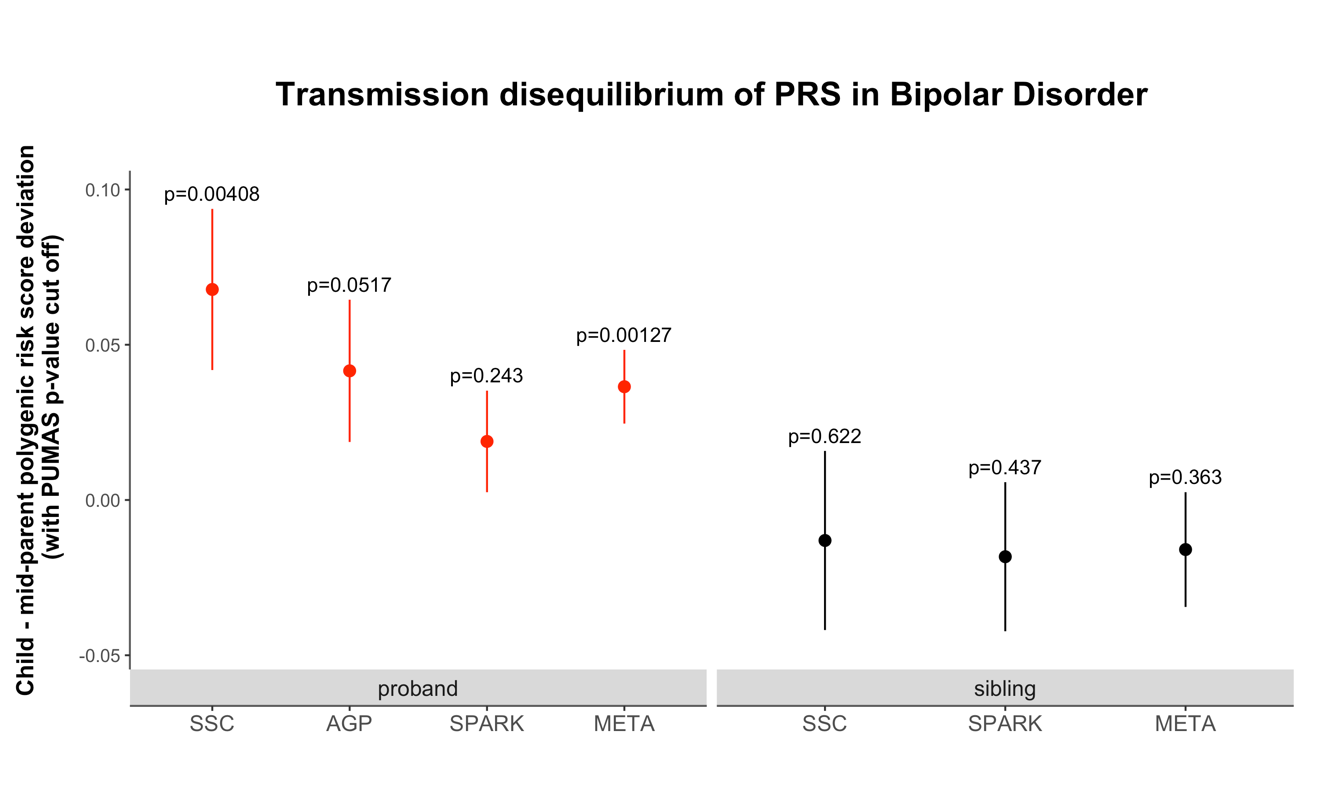

**Supplementary Figure 11. Transmission disequilibrium of PRS in different cohorts in bipolar disorder using ORIGAMI informed pTDT analysis.** Transmission disequilibrium was quantified by the ORIGAMI informed pTDT analysis with PUMAS p-value cut off. Results in probands and unaffected siblings are highlighted in different colors. The mean pTDT deviation and the SE are shown. P-values are labeled above each interval.

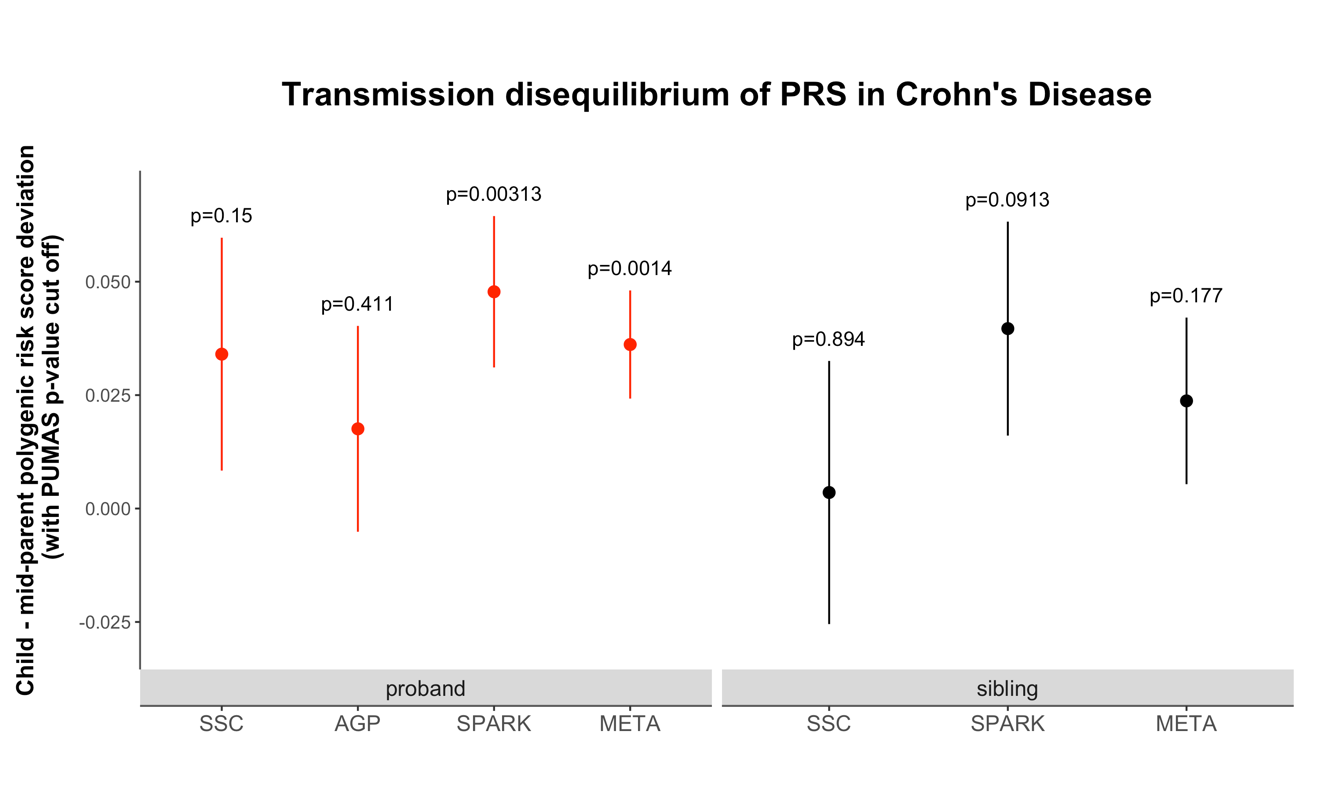

**Supplementary Figure 12. Transmission disequilibrium of PRS in different cohorts in Crohn’s disease using ORIGAMI informed pTDT analysis.** Transmission disequilibrium was quantified by the ORIGAMI informed pTDT analysis with PUMAS p-value cut off. Results in probands and unaffected siblings are highlighted in different colors. The mean pTDT deviation and the SE are shown. P-values are labeled above each interval.

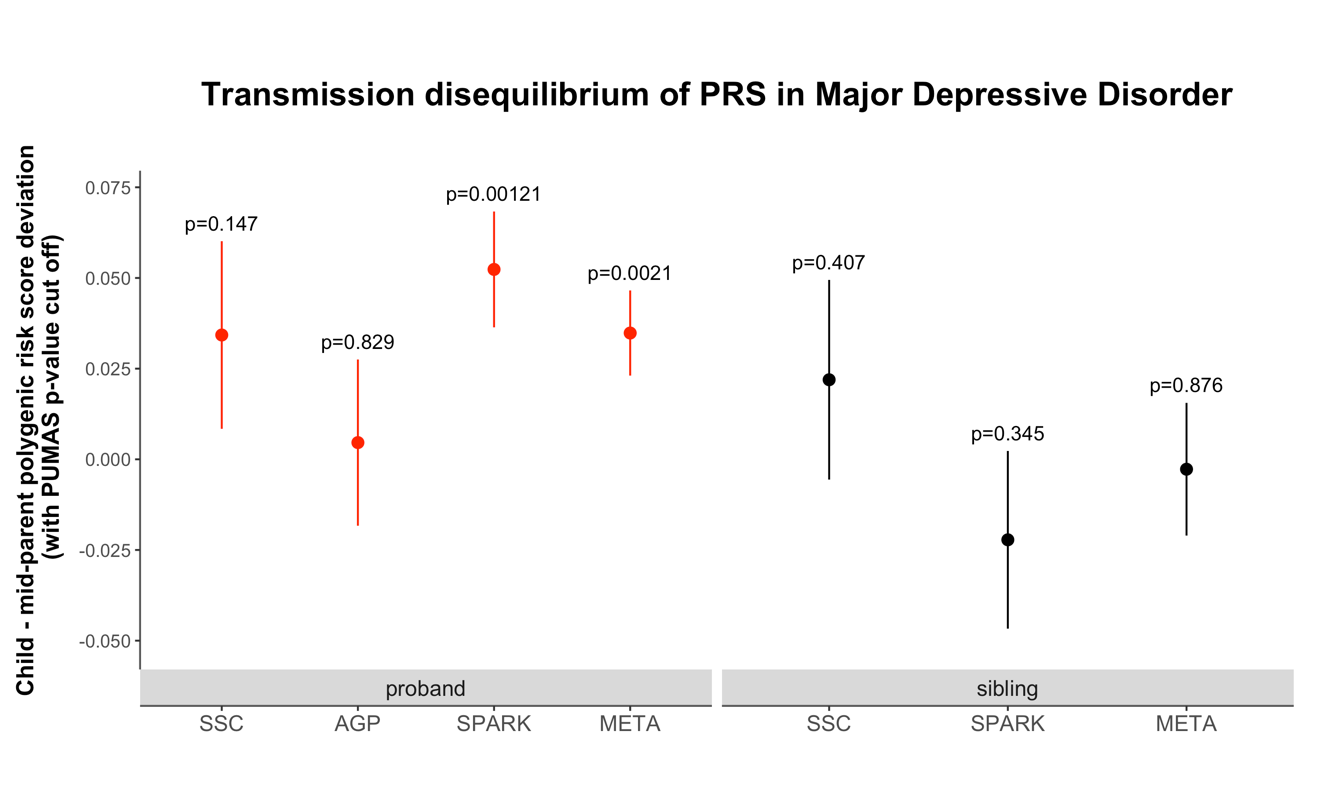

**Supplementary Figure 13. Transmission disequilibrium of PRS in different cohorts in major depressive disorder using ORIGAMI informed pTDT analysis.** Transmission disequilibrium was quantified by the ORIGAMI informed pTDT analysis with PUMAS p-value cut off. Results in probands and unaffected siblings are highlighted in different colors. The mean pTDT deviation and the SE are shown. P-values are labeled above each interval.
